# iCARE: R package to build, validate and apply absolute risk models

**DOI:** 10.1101/079954

**Authors:** Parichoy Pal Choudhury, Paige Maas, Amber Wilcox, William Wheeler, Mark Brook, David Check, Montserrat Garcia-Closas, Nilanjan Chatterjee

**Affiliations:** The Johns Hopkins University, Bloomberg School of Public Health, Department of Biostatistics, 615 N Wolfe Street, Baltimore, MD 21205.; National Cancer Institute, Division of Cancer Epidemiology and Genetics, Biostatistics Branch, 9609 Medical Center Drive, Bethesda, MD 20892-9760.; Information Management Services, Silver Spring, MD.; The Institute of Cancer Research, Division of Genetics and Epidemiology, London, United Kingdom.

**Keywords:** absolute risk, prediction, data synthesis, missing data, risk score, polygenic model, R

## Abstract

This report describes a R package, called the Individualized Coherent Absolute Risk Estimation (**iCARE**) tool, that allows researchers to build and evaluate models for absolute risk and apply them to estimate an individual’s risk of developing disease during a specified time interval based on a set of user defined input parameters. An attractive feature of the software is that it gives users flexibility to update models rapidly based on new knowledge on risk factors and tailor models to different populations by specifying three input arguments: (1) a model for relative risk, (2) an age-specific disease incidence rate, (3) the distribution of risk factors for the population of interest. The tool can handle missing information on risk factors for individuals for whom risks are to be predicted using a coherent approach where all estimates are derived from a single model after appropriate model averaging. The software allows single nucleotide polymorphisms (SNPs) to be incorporated into the model using published odds ratios and allele frequencies. The validation component of the software implements the methods for evaluation of model calibration, discrimination and risk-stratification based on independent validation datasets. We provide an illustration of the utility of **iCARE** for building, validating and applying absolute risk models using breast cancer as an example.

## 1. Introduction

Absolute risk models estimate disease risk in an upcoming time interval based on known risk factors for healthy individuals in a population, accounting for the presence of competing outcomes, such as death from other causes (Gail *et al.* 1989; Pfeiffer and Gail 2017). Absolute risk models for cancers and other diseases have important clinical and public health applications. Assessment of absolute risk of disease is fundamental for developing health intervention strategies to optimize an individual’s risks and benefits. For example, absolute risk models can be used to identify individuals who have a high risk of disease in order to target screening and disease prevention strategies (Jackson 2000; Jackson *et al.* 2005; Pharoah *et al.* 2008; Gail 2011). Decisions regarding the initiation of screening or preventive intervention are often made on the basis of age and family history, considered proxies for risk. However, there is increasing consensus in the medical community that these decisions should instead be guided directly by individualized estimates of risk, which can be obtained from absolute risk models that include a wider array of environmental and genetic risk factors. Assessment of the distribution of risks for individuals in the population allows public health researchers to weigh the risks and benefits of a given intervention, such as a screening regimen, for the entire population (Grundy 1999; Gail 2001; Murray *et al.* 2003). Absolute risk models can also be applied to assess the power of clinical trials by projecting the expected distribution of disease risk from the distribution of risk factors in a population (Gail 2011). At an individual level, absolute risk estimates can be used to counsel individuals on the basis of their personalized risk.

As large scale epidemiologic studies continue to discover new risk factors for many diseases, there is a growing demand to develop and apply models for absolute risk prediction that can facilitate translation of our understanding of etiology into tools for managing health. At present there does not exist a general software for researchers to build, update, evaluate and apply absolute risk models in R (R Development Core Team 2010), and the Individualized Coherent Absolute Risk Estimation (**iCARE**) package provides this much needed capability.

The **iCARE** package fits absolute risk models by synthesizing multiple data sources containing information on relative risks, the distribution of risk factors in the population, and age-specific incidence rates for the disease of interest and rates of competing risks. This compartmentalization allows researchers to incorporate the best available information on key model parameters, to easily update models as new information becomes available, and to tailor or extend models to particular populations. For clinical utility of absolute risk prediction models, it is critical that they are validated in independent studies that did not contribute to the model building. Model validation involves two aspects: (i) model calibration i.e., whether the model is producing unbiased estimates of risk in subjects with varying risk factor profiles; (ii) model discrimination, i.e., ability of the model to draw distinction between cases and controls. We have updated **iCARE** with a validation component, that will allow evaluation of comparative performance of models, possibly across different studies, using a unified set of methods. Releasing **iCARE** will reduce that start up time for researchers, help standardize the methodology and make it easy to share absolute risk models and make associated analyses reproducible. The package also implements methods for handling missing covariate data, which is likely to be an issue in practice, and gives special attention to the efficient incorporation of genetic factors based solely on published information.

## 2. iCARE Methodology: Synthesizing Data Sources

Here, we present the statistical framework underlying the **iCARE** package. We describe the data inputs that are required to use the tool, examples of appropriate sources for the data, and details regarding how the key inputs are used to estimate model parameters. Specifically, we explain the methodology used to estimate the baseline hazard function component of the model and the approach used to handle missing data in the risk factor profiles used in the estimation of individuals’ risks. We describe the tool’s special treatment of SNPs, which allows genetic risk factors to be incorporated into the model based on published information.

### 2.1 Model

The **iCARE** package fits a model for absolute risk, assuming the age-specific incidence rates of the disease given a set of risk factors, Z, follow the Cox proportional hazard (PH) model (Cox 1972) of the form

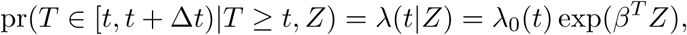

where *T* represents the time of diagnosis for the disease of interest. The model assumes that risk factors *Z* act in a multiplicative fashion on the baseline hazard function, *λ*_0_(*t*). Given this model, the absolute risk of the disease for an individual who is currently at age *a* over the time internal *a* + *τ* is defined as (Gail *et al.* 1989),

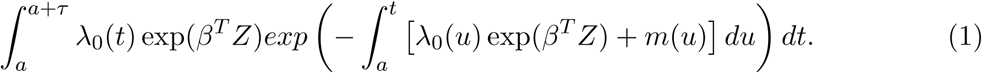

Formula (1) accounts for competing risks due to mortality from other causes through the age-specific mortality rate function *m*(*t*). In the current implementation, for simplicity, it is assumed that risk of mortality does not depend on the risk factor *Z*, but the method, in principle, can be extended to relax this assumption if covariate-specific risks of competing mortality can be estimated from external sources or models.

### 2.2 Data and Estimation

In order to build the above model and apply it for absolute risk estimation, users must provide three main data sources:

- a model for the log relative risk (or log hazard ratio) parameters: *β*
- a marginal age-specific disease incidence rate: *λ*_*m*_(*t*)
- a dataset containing risk factors for a set of representative individuals that could be used to estimate the risk factor distribution for the underlying population: *Z*_*j*_ for *j* = 1, *…, N*_*ref*_

In order to account for competing risks, an optional input with age-specific incidence rates of all-cause mortality, ideally excluding the disease of interest, *m*(*t*) should also be provided. The **iCARE** tool computes absolute risk estimates as the sum of the integrand of (1) over integer ages in the time interval of interest. The user-provided log hazard ratio parameter estimates, 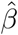, are plugged into the equation directly to carry out the computation. There are a number of ways that these input parameters may be obtained. For example, the estimates 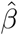 may be derived from the analysis of a prospective cohort study using a multivariate PH model. Alternatively, they may be obtained from the analysis of a case-control study using a multivariate logistic regression model adjusted for fine categories of age (Prentice *et al.* 1978). Ideally, datasets used to estimate model parameters should include information on all risk factors of interest and be large enough to provide precise estimates. When this is not available, estimates of relative risk for different risk factors could be obtained from multiple data sources (e.g., large published studies or meta-analyses). It is important that the provided estimates are adjusted for other risk factors in the model and interactions.

The second data source needed for the model is an estimate of the overall (or marginal) age-specific disease incidence rate, defined as

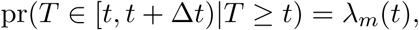

for the population of interest. This information, for example, could be available from populationbased registries, such as the United States’ Surveillance Epidemiology and End Results (SEER) cancer registry maintained by the National Cancer Institute (Howlader *et al.* 2011). Similarly, users who wish to account for competing risks must provide the optional marginal age-specific incidence rates of all-cause mortality excluding the disease of interest

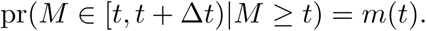

In general it is best to incorporate rates defined for fine age categories, such as 1or 5-year age strata, however **iCARE** can accommodate information on coarser age strata as well. For estimation, the age-specific disease incidence rates *λ*_*m*_(*t*) are used in combination with the third data input, a dataset of risk factors that is representative of the population of interest, to estimate the baseline hazard function, *λ*_0_(*t*).

### 2.3 Estimating the Baseline Hazard Function

Given the model of log relative risks, 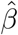, and marginal age-specific disease incidence rates, 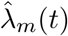, we use the following relationship to derive the baseline hazard rate

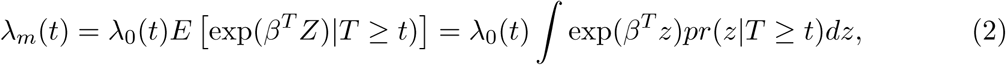

where, under the proportional hazard model,

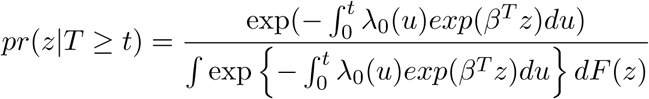

with *F* (*Z*) denoting the distribution of the risk factors in the underlying population. If the disease can be assumed to be rare, then (2) can be approximated in closed form as

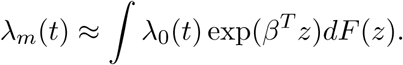

Computationally, the **iCARE** implementation starts with an initial value for *λ*_0_(*t*) based on the rare disease approximation and iterates based on formula (2) to obtain more exact estimates. This approach is closely related to an alternative formula for estimation of *λ*_0_(*t*) described by Gail *et al.* (1989). That approach involved an alternative maneuvering of the formula to allow estimation based on the risk factor distribution from a random sample of cases. In contrast, our estimation method relies on an available distribution of the risk factors for a general population. Thus, a model based on our proposed method (as implemented by **iCARE**) can be easily updated to reflect the risk factor distribution for different populations without requiring access to a sample of cases from each population of interest.

### 2.4 Specification of risk factor distribution

As described in Section 2.3, the risk factor distribution *F* (*Z*) plays a key role in calibrating the model to the marginal disease incidence rates in the underlying population. Thus, to carry out the calibration, the user must provide individual level data on the model risk factors for a sample that is representative of the underlying population. Ideally, this representative dataset would simply be the empirical distribution of *Z*, from a national survey, an epidemiologic study such as a population-based cohort or controls from a population-based case-control study sampled from the population of interest. When empirical data are available, there are no additional modeling assumptions needed. However, if complete empirical data in all risk factors are not available, users can instead provide a representative dataset that may have been simulated under modeling assumptions appropriate to the population of interest. For example, in the application illustrated later we develop a model for absolute risk of breast cancer and incorporate a representative dataset of risk factors *Z* which were simulated based on data from a combination of national surveys.

### 2.5 Handling Missing Data in Covariate Profile

In addition to providing the three data inputs for estimating model parameters, users must provide information on risk factors for the individuals to whom the model should be applied. When there is complete information for all risk factors of interest, risk estimation is as straightforward as plugging the risk factors *Z* into formula (1). However, in practice there may be missing data on some of the risk factors for individuals for whom we want to produce risk estimates.

One way to handle missing data on risk factors for a given individual is to use multiple imputation procedures (Rubin 2008). The user would obtain estimates of absolute risk using iCARE for each of the completed-by-imputation risk factor profiles for the individual, and then average the absolute risk estimates to obtain an overall estimate of the absolute risk for that individual.

The **iCARE** tool also provides an internal option for handling missing data in the covariate profile for prediction: model-free imputation based on the reference dataset of risk factors provided by the user. The methodology underlying this imputation is as follows. For any subject indexed by *i* with a covariate profile *Z*_*i*_, we define the risk score *R*_*i*_ = *β*^*T*^ *Z*_*i*_, the linear predictor associated with the user specified log relative risk model. If a subject has missing values in some of the covariates, we partition 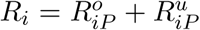, where *P* indexes the observed pattern of missing data and where 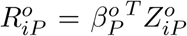 and 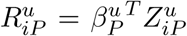 denote the corresponding “observable” and “unobservable” components of the risk score. In general, this partitioning depends on which columns of the design matrix of the original model can be specified by the observed set of covariates for a given individual’s risk factor profile. Given this partitioning, the absolute risk, *AR*, of the individual *i* is defined by

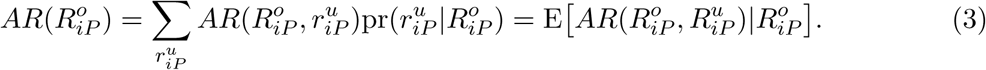

The absolute risk for the *i*-th individual is obtained by averaging over possible values for the unobserved component of the risk score given the value of the observed component of the risk score. As all the risk scores are scalar quantities, one can estimate the conditional distributions 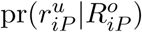 in a fairly non-parametric fashion using the user-specified reference dataset.

In particular, to carry out (3) for a given covariate profile with missing data, the method finds subjects in the reference dataset that are similar on the basis of the observable component of the risk score, 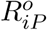, and take as the risk estimate the average of the full model risk, 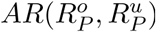, for the referent subjects identified to be similar. Specifically, the observable risk scores 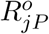 are obtained for *j* = 1,..*N*_*ref*_ in the reference dataset, categorized into single percentile strata, and the individual’s 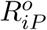 is matched to one of the strata. The reported risk for the individual is then computed by averaging over the values of the full 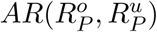 for all referent subjects in this matching stratum. This method can be viewed as a type of “hot deck” imputation based on the risk score, which is popular in survey literature.

### 2.6 Special option for SNP markers

As large genome-wide association studies continue to discover low penetrant, common SNPs associated with risk of complex chronic diseases, it is important to investigate the utility of the SNPs, in combination with other risk factors, for public health strategies of disease prevention. Evaluation of absolute risk, as opposed to relative risk which is typically used for summarizing associations, is fundamental for these public health applications. Due to the importance of SNP markers in absolute risk models and natural assumptions specific to genetic data, the **iCARE** package provides a number of options for incorporating SNPs into the model.

Users can include individual SNPs in the model, or include a polygenic risk score (PRS), in the same way as any other risk factor as long as all input components can be identified. This allows researchers to specify interactions between SNPs and other risk factors in the model or to include PRSs with more complex weighting structures, if desired. However, to include SNPs this way, a reference dataset must be provided that has the individual SNPs (or the PRSs) for all subjects. If the reference dataset is not available, the researchers may need to estimate this reference distribution by creating a simulated dataset of individuals who are representative of the underlying population.

Alternatively, the **iCARE** package also provides a special approach for handling independent SNPs, which requires that the user only provide information on the odds ratio *θ*_*k*_ and population allele frequency *f*_*k*_ for each SNP to be included. **iCARE** internally creates a PRS from all provided SNPs weighted by the odds ratios,

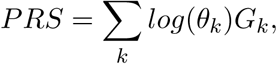

where *G*_*k*_ denotes the SNP genotype status of individuals, coded as the number of non-referent alleles they carry (with respect to the referent allele for which the odds ratios are reported). In general, iCARE assumes this PRS to be distributed independently of all other covariates. However, if a family history (fh) variable is included in the model, then the method allows a simple adjustment to account for correlation between PRS and family history. The adjustment method assumes the latter is coded as a binary indicator of presence or absence of disease among first-degree relatives. In particular, when the model risk factors include family history, **iCARE** provides the option to adjust the log odds ratio associated with family history using the formula:

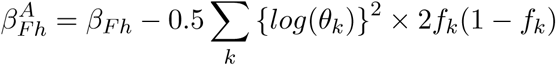

with *θ*_*k*_ denoting the disease odds ratio of the SNPs, unadjusted for family history. This adjustment reflects the fact that, with the addition of SNPs into the model, the effect of family history is attenuated by a magnitude that is proportional to the degree of heritability explained by the SNPs. This treatment should be applied only when the provided *β*_*F* *h*_ represents the effect of a binary variable for first-degree family history, unadjusted for the SNPs.

Users may provide relative risk estimates for family history that are already adjusted for the SNPs in the model, and if so they should simply not select the option for the family history adjustment.

One important way in which this approach treats SNPs differently involves the reference dataset of risk factors. This dataset might come from a national survey; however, a national survey is unlikely to have genotyped individuals, particularly for the exact set of SNPs to be included in the model. Recognizing this, for user convenience, the reference dataset just needs to include non-genetic risk factors and **iCARE** will simulate SNP genotype values based on the provided allele frequencies for the population. The SNPs are multiply imputed with user-specified number of imputations (`n.imps`) for each subject in the reference dataset. The method assumes that the SNPs are independent and that the genotype distributions follow Hardy-Weinberg Equilibrium. Specifically, the joint distribution of SNP genotypes and other risk factors (*X*) are assumed to follow the decomposition

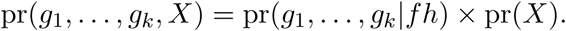

If family history of the disease is included in the model as a binary risk factor indicating the presence or absence of any first-degree relative with disease history, assuming that the disease is rare, we approximate

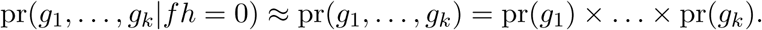

The distribution of SNP genotypes among subjects with family history is approximated as

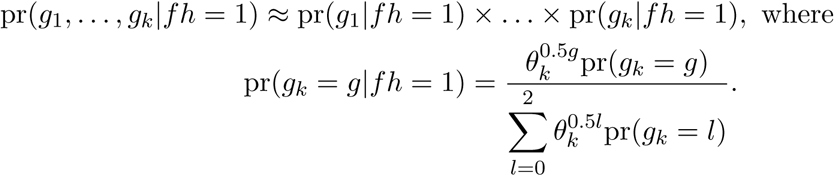

The above approximation is derived under the assumption of rare disease and multiplicative effect of SNPs on the risk of the disease. If family history is not indicated to be in the model, and is thus not provided for each reference dataset subject, we impute the SNPs based on the unconditional distribution for independent SNPs in Hardy-Weinberg equilibrium.

It is possible that SNP information may also be missing in the covariate profiles for whom the model will be applied to estimate risk. In this case, SNPs are treated the same as all other risk factors and handled according to the methodology given in Section 2.5. This approach is equivalent to averaging over the possible values of the missing SNPs according to the population distribution, taking advantage of any known SNPs in the genotype profile.

### 2.7 Model validation methods

Suppose we are given a model to predict the absolute risk of a disease. In other words, the different pieces of information needed to estimate absolute risk in (1) are given to us. Moreover, we also have an independent prospective cohort study (or a case-control study nested within a cohort), that we would like to use to validate the absolute risk model. Given below are the steps for implementing the model validation methods in the independent study.

#### Model calibration

Model calibration attempts to answer the question whether the risk prediction model is producing unbiased estimate of risk for subjects with varying risk factor profiles. The observed followup (*T*_*F*_) for a subject in the validation cohort is defined as the time from study entry to the last contact or linkage for acquiring information. Often, researchers desire to validate model for absolute risk over a fixed year (*τ*) of follow-up (e.g., 5-year or 10-year). For calculation of both observed and expected risks, the subjects, who did not develop the disease by *τ* years and were followed up longer than *τ* years, were censored at *τ* years i.e., for model calibration their absolute risk was estimated over *τ* years. For all other subjects the absolute risk is estimated over the minimum of *τ* years and observed followup. Figure 1 provides an illustration. The subjects, who developed the disease by *τ* years, were considered as cases and those, who did not develop the disease by *τ* years, were designated as controls. The overall *τ* -year observed risk can be computed as the observed proportion of cases in the study. For each subject, the predicted *τ* -year risk can be supplied by the user. Alternatively, it can be computed by **iCARE**, using equation (1), with the input components supplied by the user. It is also possible to validate model over the entire follow-up period of study instead of pre-specifying a fixed follow-up year.

**Figure 1:**
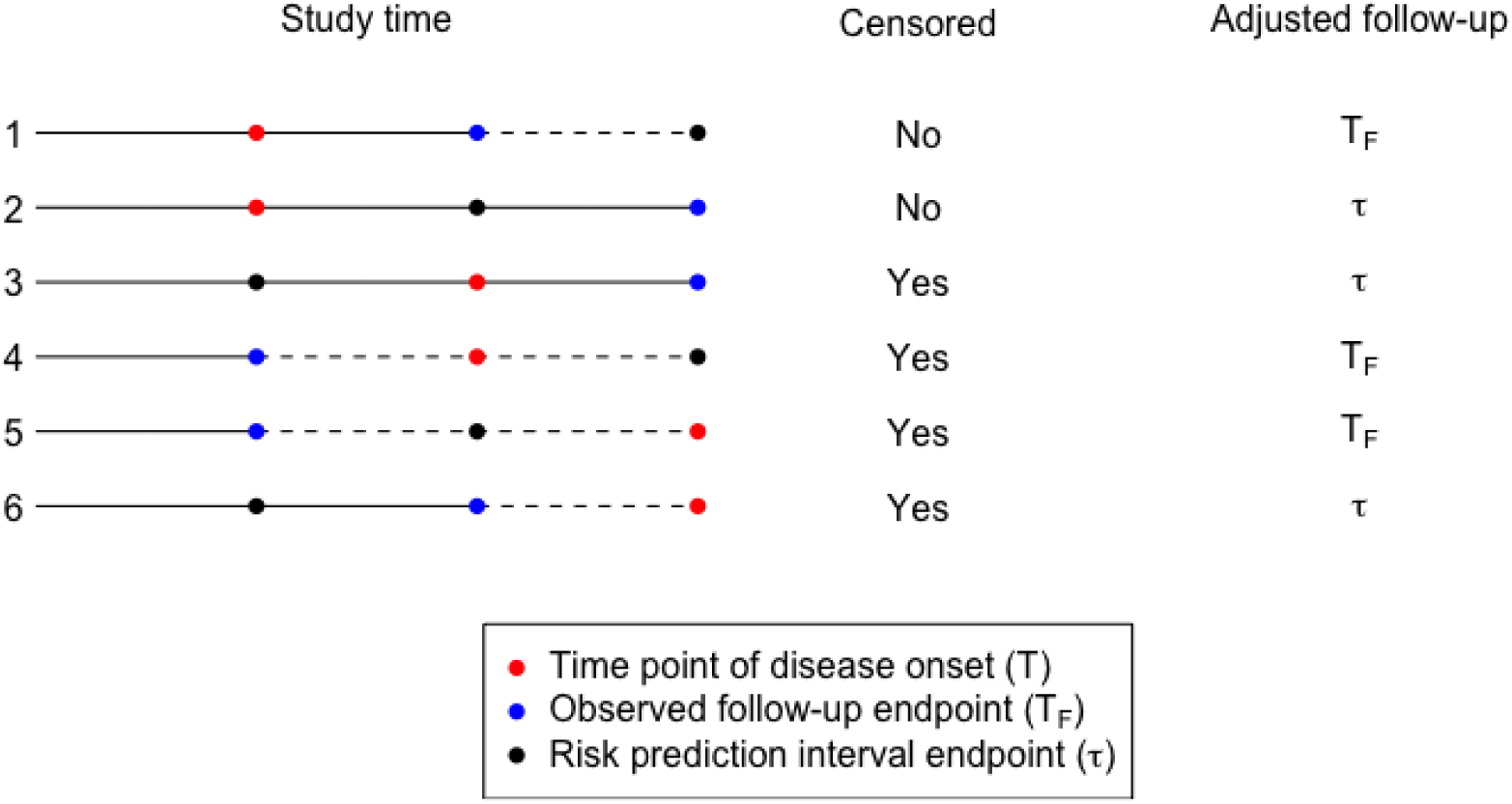
Illustration of calculation of adjusted followup for the different kinds of subjects in independent validation of a risk prediction model: *T* := time (in years) since study entry to onset of disease; *T*_*F*_ := observed follow-up (in years) since study entry; *τ* := length of risk prediction interval (in years) starting at study entry; the black solid line indicates that the subject is being followed up and the black dotted line indicates that the subject is lost to follow-up.

We categorize the validation cohort based on low risk to high risk of disease based on a prespecified number of categories (e.g., deciles) of the risk score *R*_*i*_, defined earlier in section 2.5. We denote the categorical version of risk score as 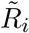. The categories are indexed by *c* = 1, …, *C* (e.g., C = 10 for deciles). At each category, we compute the observed risk, 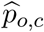, defined as the observed proportion of cases in that category. For comparison, we also compute an average of the subject specific predicted absolute risk of disease, 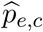, over the subjects in that category. An influence function based variance formula for 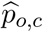 is implemented (details in Appendix). The 95% Wald based confidence intervals for 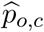 are computed using this variance formula. A Hosmer-Lemeshow type test for goodness of fit is also implemented, where the test statistic has the form:

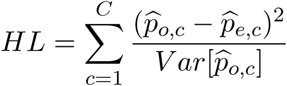

The above HL statistic approximately follows a chisquare distribution with *C* degrees of freedom.

The software also has capabilities to perform calibration on the scale of relative risks. The observed relative risk in each category is defined as the ratio of the observed absolute risk in that category and the overall observed absolute risk in the validation study. For comparison, the predicted relative risk is computed as the average of the predicted absolute risks over the subjects in that category and the overall average of the predicted absolute risks in the validation study. A variance formula for the observed relative risk is implemented using delta method (details in Appendix) and is used to compute the 95% Wald based confidence intervals. A chisquare goodness of fit test for relative risk is also implemented.

In a cohort study, data on some expensive biomarkers (e.g., genetic risk factors) are often collected only on a judiciously selected sub-sample nested within the original cohort (e.g., a nested case-control sample). In this case, the estimates of absolute and relative risks need to be adjusted to account for the non-random sampling using sampling weights. An inverse-probability weighted (IPW) approach is implemented to estimate the absolute risks. Let *I*_*i*_denote the indicator of inclusion of a cohort subject (indexed by *i*) to the nested case-control sample and *Y*_*i*_ denote the case-control status. Denote by *π*_*i*_(*Y*_*i*_, *Z*_*i*_) = *P* [*I*_*i*_ = 1|*Y*_*i*_, *Z*_*i*_], the sampling weight, i.e., the probability of inclusion for a subject indexed by *i*. The cut-points for the risk score categories are calculated using these sampling weights by inverting a weighted empirical distribution function of the risk score. The adjusted estimates of observed and expected risks, overall and at each category, are computed as the corresponding weighted averages (instead of simple averages) with weights being inverse of the sampling weights. A variance estimator of the observed risk in each category is also implemented using the influence function approach (details in Appendix). The 95% Wald based confidence interval is computed using this variance formula. The Hosmer-Lemeshow type goodness of fit test statistic also uses this variance formula in its denominator.

#### Model discrimination

For well calibrated models, it is critical that the risk factors used in the model have good discriminatory power i.e., ability to differentiate between cases and controls. A risk prediction model with good discriminatory ability ensures that there is a good amount of spread in the risk distribution of the subjects to facilitate risk stratification leading to better evaluation of risks and benefits. A common measure of model discrimination is the Area Under the Curve (AUC) defined as *δ* = *P* [*R*_1_ *> R*_0_], i.e., the probability that the risk score for a case (*R*_1_) is higher than the risk score for a control (*R*_0_). For a cohort study, this probability can be estimated using the empirical proportion of case-control pairs for which the risk score of the case is greater than that of the control. For this setting, the asymptotic variance formula of this estimator (DeLong *et al.* 1988) is implemented and used to compute the 95% Wald based confidence interval. For a nested case-control study, we implement an inverse probability weighted estimator of AUC [(Cai and Zheng 2011, 2012; Zheng *et al.* 2013; Zhou *et al.* 2013; Yao *et al.* 2015)]. We also implement an influence function based variance estimate. The influence function representation and the variance formula are shown in the Appendix and further details are provided in Pal Choudhury *et al.* (2017). The 95% Wald based confidence interval of this estimator is computed using this variance estimate.

## 3. Existing R packages

In this section, we describe some of the unique aspects of iCARE compared to existing R packages for building and validating a model for predicting risk of a disease (see Table 1 for a summary). There are additional software packages developed in other platforms that allow risk calculation of specific cancers, e.g., **IBIS** Breast Cancer Risk Evaluation Tool (Tyrer *et al.* 2004; Cuzick 2017) and Breast and Ovarian Analysis of Disease Incidence and Carrier Estimation Algorithm **(BOADICEA)** (Antoniou *et al.* 2004, 2008; Lee *et al.* 2014; Cunningham and Antoniou 2018).

**Table 1:** Summary of features of existing R packages for risk prediction. ✓ denotes the presence of the feature and *×* denotes absence of the feature.

| Packages | Model building |  |  |  | Model validation |  |  |
| --- | --- | --- | --- | --- | --- | --- | --- |
|  | Calibration to population incidence | Detailed family history | Special option for SNP markers | Imputation of missing risk-factors | Full cohort | Two-phase study | Imputation of missing risk-factors |
| **riskRegression**<sup>1</sup> | × | × | × | × | ✓ | × | × |
| **predictABEL**<sup>1</sup> | × | × | × | × | ✓ | × | × |
| **BCRA**<sup>2</sup> | ✓<sup>3</sup> | × | × | × | × | × | × |
| **BayesMendel**<sup>2</sup> | × | ✓ | × | ✓ | × | × | ×<sup>6</sup> |
| **rmap** | × | × | × | × | ✓ | ✓<sup>5</sup> | × |
| **iCARE**<sup>2</sup> | ✓<sup>3</sup> | × | ✓<sup>4</sup> | ✓ | ✓ | ✓<sup>5</sup> | ✓<sup>6</sup> |

<sup>1</sup> These packages include some functions for model building (see Section 3.1), but those approaches do not demonstrate the key features shown in the above table.
<sup>2</sup> Capability to use information from multiple data sources, e.g., **BCRA** and **iCARE** can use relative risk parameters from cohort or case-control studies and disease incidence and mortality rates from population registries.
<sup>3</sup> **BCRA** estimates baseline hazard and calibrates the model to the underlying population incidence rates using distribution of risk-factors from cases in a specific study that may not be representative of the general population. This step is implemented in **iCARE** using a reference dataset that provides information on the distribution of risk-factors in the general population.
<sup>4</sup> **iCARE** includes the special option in which independent SNP markers can be included using published estimates of odds ratios and allele frequencies.
<sup>5</sup> Inverse probability weighted estimators of model validation statistics are implemented, accounting for bias due to non-random sampling using sampling weights.
<sup>6</sup> **BayesMendel** incorporates imputation methods for certain risk-factors (e.g., age), but they do not implement any method of validating risk prediction models. **iCARE** implements an inbuilt imputation approach to deal with missing risk-factors using a reference risk-factor dataset representative of the underlying population. The standardized model validation methods implemented in **iCARE** can take advantage of this inbuilt feature to impute missing risk-factors in the validation study.

### 3.1 Model building

The **riskRegression** package (Gerds *et al.* 2017, 2012; Benichou and Gail 1990) implements various regression based methods for estimation of cumulative incidence in a cohort study setting with time-to-event outcome subject to censoring. The methods cannot be easily adapted to use information from multiple data sources, such as relative risk parameters from case-control studies and disease incidence information from population registries. In the **PredictABEL** package (Kundu *et al.* 2015), the conditional probability of disease given risk factors is estimated using a logistic regression model. The target quantity is different from absolute risk. The **BayesMendel** package (Chen *et al.* 2004) implements Bayesian approaches to genetic risk prediction using information from detailed family history, genetic test results and some tumor characteristics. It is based on Mendelian inheritance mechanisms in the setting of family based studies of some specific cancers (e.g., breast cancer, colorectal cancer, pancreatic cancer, endometrial cancer). As opposed to their method, we focus on risk prediction in the general population using information from common genetic variants and epidemiologic risk factors in settings where detailed family history may not be available.

The **BCRA** package (Zhang 2018) is developed to calculate risk projections for invasive breast cancer based on the Gail model (Gail *et al.* 1989), and its subsequent extensions in the different ethnic groups (Costantino *et al.* 1999; Gail *et al.* 2007; Matsuno *et al.* 2011; Banegas *et al.* 2016). The model uses information from some epidemiologic risk factors and family history of breast cancer. The basic setup of **iCARE** and **BCRA** are similar as both build models for absolute risk using information from different data sources and they use the same formula for the absolute risk based on an underlying Cox proportional hazard model (Gail *et al.* 1989). There are, however, important differences in the underlying methodology. For estimating baseline hazard and calibrating the model to the underlying population incidence rates, **BCRA** uses the distribution of risk factors in cases of the study population. In contrast, we perform this calibration step using the distribution of risk factors coming from the general population. Our approach allows for developing risk models that reflect the distribution of risk factors in the underlying population without having to use the distribution of risk factors from cases of a specific study that may not be representative of the general population. While the **BCRA** does not have an inbuilt machinery to handle missing risk-factor data, the **iCARE** uses a coherant approach that involves internal imputation of missing risk-factors from the same model with appropriate model averaging.

### 3.2 Model validation

The **riskRegression** and **PredictABEL** packages have limited capability of model validation in a cohort study setting. The **rmap** package (Gong 2016; Gong *et al.* 2014; Whittemore and Halpern 2016) implements methods for independent evaluation of risk prediction models in studies that may involve two phase designs. One unique feature of **iCARE**, that is not available in other packages, is that it includes an exhaustive set of functions so that model building and model validation can be implemented in tandem. Thus the **iCARE** based model validation procedures can take advantage of the inbuilt features of the package to deal with summary level genetic information and imputation of missing risk factors.

## 4. Using the iCARE package

In this section, we demonstrate how to use **iCARE** to build and apply two absolute risk models for breast cancer: one with only SNP markers, and one with risk factors and SNPs. We also illustrate the use of the validation component of the package. The results of these illustrative examples were generated using the RStudio platform (Team *et al.* 2018)

The main function to compute absolute risk in **iCARE** is `computeAbsoluteRisk`. The input arguments to this function are named with the prefix “`model.`” or “`apply.`” according to whether they are used primarily for model building or application respectively.

To begin, the R package and the example dataset object `bc_data` should be loaded:

~~~
R> library(“iCARE”)
R> data(“bc_data”, package=“iCARE”)
R> set.seed(50)
~~~

### Example 1: SNP-only Model

To specify a SNP-only model, we must input the marginal age-specific disease incidence rates of breast cancer, `bc_inc`, and the SNP information matrix, `bc_15_snps`, that has three columns named: `snp.name`, `snp.odds.ratio`, and `snp.freq`. Marginal age-specific incidence rates of competing risks (`mort_inc`) are optional. We include them in this example.

Here, `bc_15_snps` contains published information on odds-ratios and allele frequencies of 15 SNPs identified to be associated with breast cancer risk by a recent genome-wide association study (Michailidou *et al.* 2015). `bc_inc` contains age-specific incidence rates of breast cancer from Surveillance, Epidemiology and End Results (SEER) Program, and `mort_inc` has age-specific incidence rates of all-cause mortality from the WONDER mortality database (National Center for Health Statistics (NCHS) 2014). In fitting a SNP-only model, the reference dataset need not be provided as **iCARE** will impute the reference SNP distribution based on SNP allele frequencies. The function call below builds an absolute risk model based on 15 SNPs for breast cancer and applies the model to estimate risk of breast cancer in the interval from age 50 to age 80:

~~~
R> res_snps_miss = computeAbsoluteRisk(model.snp.info = bc_15_snps,
                      model.disease.incidence.rates = bc_inc,
                      model.competing.incidence.rates = mort_inc,
                      apply.age.start = 50, apply.age.interval.length = 30,
                      return.refs.risk = TRUE)
~~~

For this SNP-only model, we first opted for not providing any new profiles for estimation (i.e., no `apply.snp.profile` input). In this case, **iCARE** simulates 10,000 SNP profiles internally for the reference dataset and reports as the risk estimate the average of the risks estimated from the profiles: 0.09583. We can access the estimated risks for the (simulated) referent profiles and obtain summary information by calling:

~~~
R> summary(res_snps_miss$refs.risk)
Risk_Estimate
Min. :0.07652
1st Qu.:0.09198
Median :0.09575
Mean :0.09583
3rd Qu.:0.09950
Max. :0.11966
~~~

From this, we learn that on average women of age 50 have a 9.58% chance of being diagnosed with breast cancer before age 80, and that the 15-SNP model stratifies breast cancer risk from a minimum risk of 7.65% to a maximum risk of 11.97% in the age interval 50 years to 80 years.

Next, suppose we want to predict risk for three specific women whom we have genotyped; we can then call:

~~~
R> res_snps_dat = computeAbsoluteRisk(model.snp.info = bc_15_snps,
                      model.disease.incidence.rates = bc_inc,
                      model.competing.incidence.rates = mort_inc,
                      apply.age.start = 50, apply.age.interval.length = 30,
                      apply.snp.profile = new_snp_prof,
                      return.refs.risk = TRUE)
~~~

Now our output `res_snps_dat$risk` contains the risk estimates for the three women whose genotype profiles we provided. Additionally, we have put the option: `return.refs.risk = TRUE`. Hence, `res_snps_dat$refs.risk` returns the risk estimates for the reference dataset based on 10,000 internal simulations. These results allow us to create a useful plot, like Figure 2, showing the distribution of risks in our reference dataset and to add the risks of the three women to see where they fall on the population distribution. We use the code:

~~~
R> plot(density(res_snps_dat$refs.risk),
              xlim = c(0.07,0.14), xlab = “Absolute Risk of Breast Cancer”,
              main = “Referent SNP-only Risk Distribution: Ages 50-80 years”)
R> abline(v = res_snps_dat$risk, col = “red”)
R> legend(“topright”, legend = “New profiles”, col = “red”, lwd = 1)
~~~

**Figure 2:**
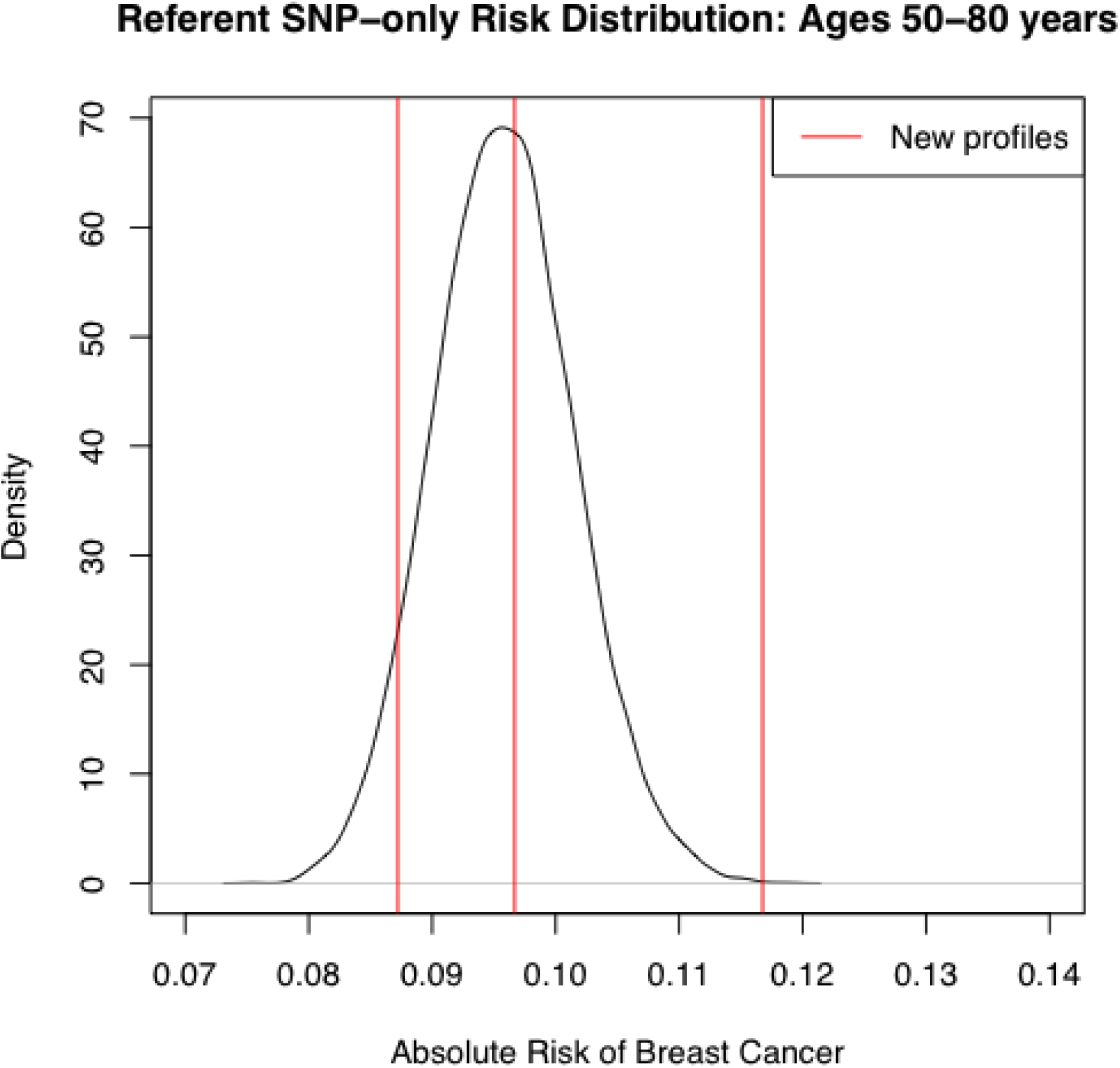
Estimated absolute risk of three women overlaid on the population distribution of Risk in the age interval: 50-80 years.

In this example, the first genotype profile had missing data on two SNPs. This demonstrates the capability of **iCARE** to produce risk estimates when there is missing data in the risk factor profile without compromising on user convenience.

### Example 2: Breast Cancer Risk Model with Risk Factors and SNPs

In this example, we illustrate the use of iCARE to build models of absolute risk with both epidemiologic risk factors and SNPs. For illustrative purpose, we use the risk factors family history and parity. Family history is the binary indicator of presence/absence of disease in first degree relatives. Parity is defined as the number of child births. The first step is to prepare the input parameters `bc_model_cov_info`, containing the epidemiologic risk factor information and `bc_model_formula`, containing the model formula object for the part of the model with epidemiologic risk factors only:

~~~
R> v1 = list(); v1$name = “famhist”; v1$type = “continuous”
R> v2 = list(); v2$name = “parity”; v2$type = “factor”;
                                                          v2$levels = c(0,1,2,3,4)
R> bc_model_cov_info = list(v1,v2)
R> bc_model_formula = observed.outcome ˜ famhist + as.factor(parity)
~~~

Having prepared the data sources, we can now run:

~~~
R> res_covs_snps = computeAbsoluteRisk(model.formula = bc_model_formula,
                       model.cov.info = bc_model_cov_info,
                       model.snp.info = bc_15_snps,
                       model.log.RR = bc_model_log_or,
                       model.ref.dataset = ref_cov_dat,
                       model.disease.incidence.rates = bc_inc,
                       model.competing.incidence.rates = mort_inc,
                       model.bin.fh.name = “famhist”,
                       apply.age.start = 50,
                       apply.age.interval.length = 30,
                       apply.cov.profile = new_cov_prof,
                       apply.snp.profile = new_snp_prof,
                       return.refs.risk = TRUE)
~~~

We indicate `model.bin.fh.name = “famhist”` to allow the software to properly attenuate the log odds ratio for family history to account for the addition of the 15 SNPs. With the exception of `model.bin.fh.name`, which is always optional, all arguments listed in `green` should either be included or excluded in the function call as a set. This is to say that if one is included, then all should be included.

The above function call fits an absolute risk model with risk factors family history and parity additively with the 15 SNPs associated with breast cancer. In a model that includes risk factors, such as this one, we must supply the model formula, the risk factor information, the log odds ratios for the risk factors, and a reference dataset of risk factors to build the model. The `model.cov.info` input tells the function that family history can be treated as a continuous variable, though it only has levels 0 and 1, and that parity should be treated as a factor variable with levels 0,1,2,3,4 indicating the number of childbirths for a given subject. Here, the `bc_model_log_or` input contains the log odds ratios for family history and parity, from a logistic regression model adjusted for cohort and fine categories of age in the Breast and Prostate Cancer Cohort Consortium (Campa *et al.* 2011; Joshi *et al.* 2014). The `ref_cov_dat` dataset was created by simulation from the National Health Interview Survey (NHIS) and the National Health and Nutrition Examination Survey (NHANES), which are representative of the US population.

In addition to summarizing and plotting the risk estimates, **iCARE** includes an option to view more detailed output, by calling:

~~~
R> print(res_covs_snps$details)
~~~

This reports the interval start and end ages over which absolute risk was computed, the entire covariate profile to which the model was applied (SNPs and risk factors, if applicable), and the resulting risk estimate, as shown below.

|  | Int_Start | Int_End | Risk_Estimate | rs12405132 | rs12048493 | rs72755295 | rs6796502 |
| --- | --- | --- | --- | --- | --- | --- | --- |
| 193502 | 50 | 80 | 0.08077312 | NA | NA | 0 | 0 |
| 126252 | 50 | 80 | 0.07709955 | 0 | 0 | 0 | 0 |
| 15756 | 50 | 80 | 0.12955390 | 0 | 1 | 0 | 1 |
|  | rs13162653 | rs2012709 | rs7707921 | rs9257408 | rs4593472 | rs13365225 | rs13267382 |
| 193502 | 1 | 1 | 0 | 1 | 1 | 1 | 0 |
| 126252 | 2 | 1 | 0 | 1 | 1 | 0 | 1 |
| 15756 | 0 | 1 | 0 | 1 | 1 | 0 | 1 |
|  | rs11627032 | rs146699004 | rs745570 | rs6507583 | famhist | parity |  |
| 193502 | 0 | 1 | 2 | 0 | 0 | 2 |  |
| 126252 | 1 | 0 | 1 | 0 | 0 | 4 |  |
| 15756 | 0 | 0 | 0 | 0 | 1 | 4 |  |

### Illustration of Validation Component

We illustrate the working of the model validation component of **iCARE** by evaluating a risk prediction model with both epidemiologic risk factors and SNPs using data simulated using information from the Prostate, Lung, Colorectal and Ovarian (PLCO) Cancer Screening Trial. To be more specific, the risk factors included are family history, parity and the 15 SNPs from the earlier example.

The package includes an example of full cohort in `validation.cohort.data`. There are 50,000 subjects with information on age of study entry, age of study exit, time to disease onset, disease status and epidemiologic risk factors (i.e., family history and parity). A common scenario in modern cohort studies is that the genetic factors are not measured on all the subjects in the cohort, but only on a judiciously selected sub-sample. The package also includes a casecontrol sample (`validation.nested.case.control.data`) of 2694 women nested within the full cohort with information on age of study entry, age of study exit, time to disease onset, disease status, epidemiologic risk factors and 15 breast cancer associated SNPs. Further details on how the full cohort and nested case-control data were simulated are provided in the Appendix.

To validate a model with epidemiologic risk factors and SNPs, we will use the nested casecontrol sample. The calibration and discrimination statistics need to be adjusted for the non-random sampling using sampling weights (i.e., probability of selection of a subject to the nested case-control sample). When sampling weights are not provided, we recommend using post-hoc estimation by assuming parametric models for selection mechanism. We illustrate the sampling weight calculation through a logistic regression model of inclusion on disease status, age of study entry, observed follow-up and interaction of disease status with age of study entry and observed follow-up.

The first step is to identify the subjects in the full cohort who are included in the nested sub-sample through matching of subject identifiers and create the indicator of inclusion:

~~~
R> validation.cohort.data$inclusion = 0
R> subjects_included = intersect(validation.cohort.data$id,
                               validation.nested.case.control.data$id)
R> validation.cohort.data$inclusion[subjects_included] = 1
~~~

The next step is to fit a logistic regression model of inclusion depending on the case/control status, age of study entry and observed followup using the R function `glm`, as shown below:

~~~
R> validation.cohort.data$observed.followup =
                           validation.cohort.data$study.exit.age –
                                  validation.cohort.data$study.entry.age
R> selection.model = glm(inclusion ˜ observed.outcome
                          * (study.entry.age + observed.followup),
                                    data = validation.cohort.data,
                                    family = binomial(link = “logit”))
R> validation.nested.case.control.data$sampling.weights =
      selection.model$fitted.values[validation.cohort.data$inclusion == 1]
~~~

In the above code chunk, the line highlighted in blue shows the call to the `glm` function to fit the selection model. Once we have the sampling weights, we can call the `ModelValidation` function in **iCARE** that implements the validation analysis. We specify the risk prediction model formula with epidemiologic risk factors only in `bc_model_formula`. The list object `risk.model` combines all the input parameters required by the `computeAbsoluteRisk()` function, used internally by the `ModelValidation` function to estimate absolute risks in the validation study.

~~~
R> bc_model_formula = observed.outcome ˜ famhist + as.factor(parity)
R> data = validation.nested.case.control.data

R> risk.model = list(model.formula = bc_model_formula,
                       model.cov.info = bc_model_cov_info,
                       model.snp.info = bc_15_snps,
                       model.log.RR = bc_model_log_or,
                       model.ref.dataset = ref_cov_dat,
                       model.ref.dataset.weights = NULL,
                       model.disease.incidence.rates = bc_inc,
                       model.competing.incidence.rates = mort_inc,
                       model.bin.fh.name = “famhist”,
                       apply.cov.profile = data[,all.vars(bc_model_formula)[-1]],
                       apply.snp.profile = data[,bc_15_snps$snp.name],
                       n.imp = 5, use.c.code = 1, return.lp = TRUE,
                       return.refs.risk = TRUE)
R> output = ModelValidation(study.data = data,
                           total.followup.validation = TRUE,
                           predicted.risk.interval = NULL,
                           iCARE.model.object = risk.model,
                           number.of.percentiles = 10)
~~~

In the above function call, the argument marked in blue is the list of input parameters that are used by iCARE to build an absolute risk model and compute model based estimates of risk in the validation study. The arguments `study.data` inputs the nested case-control sample used for model validation. In this example, we have set `total.followup.validation = TRUE` and `predicted.risk.interval = NULL` to indicate that we are validating the risk over the duration of follow-up of the subjects in the validation study. One can set these parameters appropriately to validate *τ* -year risk in a given study. We have set `number.of.percentiles= 10` to indicate that the categories of low risk to high risk of disease were defined based on the deciles of the risk score.

The above function call displays the following information on the R console:

~~~
[1] “Dataset: Example Dataset”
[1] “Model Name: Example Risk Prediction Model”
[1] “Model Formula: observed.outcome ˜ famhist + as.factor(parity)”
[1] “Risk Prediction Interval: Observed Followup”

[1] “Number of study subjects: 2694”

[1] “Number of cases: 1112”

[1] “Follow-up time (years) [mean,range]: [ 9.782, (5, 13) ]”

[1] “Baseline age (years) [mean,range]: [ 62.721, (50, 72) ]”

[1] “Absolute Risk Calibration”

             Hosmer and Lemeshow goodness of fit (GOF) test for Absolute Risk

data: observed.frequency, expected.frequency
Chisquare = 11.086, df = 10, p-value = 0.3509

[1] “Relative Risk Calibration”

             Goodness of fit (GOF) test for Relative Risk

data: observed.frequency, expected.frequency
Chisquare = 10.685, df = 9, p-value = 0.2979

[1] “Model Discrimination”
[1] “Estimate of AUC: 0.54”
[1] “95% CI of AUC: (0.517, 0.563)”

[1] “Overall Expected to Observed Ratio”
[1] “Estimate: 1.024”
[1] “95% CI: (0.948, 1.107)”
~~~

The `ModelValidation` function returns additional information about the results of model validation as a list after function execution. Details on this output object are available in the help pages of the package. The `plotModelValidation` can be used create a useful plot showing the absolute risk calibration, relative risk calibration, distribution of risk scores in cases and controls and study specific disease incidence rates overlaid on the population incidence rates. A typical call to this function is as follows:

~~~
R> plotModelValidation(study.data = data,validation.results = output)
~~~

The first input parameter is the validation study data and the second input parameter is a list of all the results returned by the `ModelValidation` function. The above function call produces the plot shown in Figure 3. There are additional optional parameters that the user can specify to tailor the display of the plots (e.g., amount of smoothing of the density plots). There could be alternative scenarios where the predicted risk and risk scores for the validation subjects are already available. In that case `ModelValidation` function has options to skip the model building and use the user supplied predicted risk and risk scores to implement model validation methods. For further details, the user is recommended to consult the help pages available with the package.

**Figure 3:**
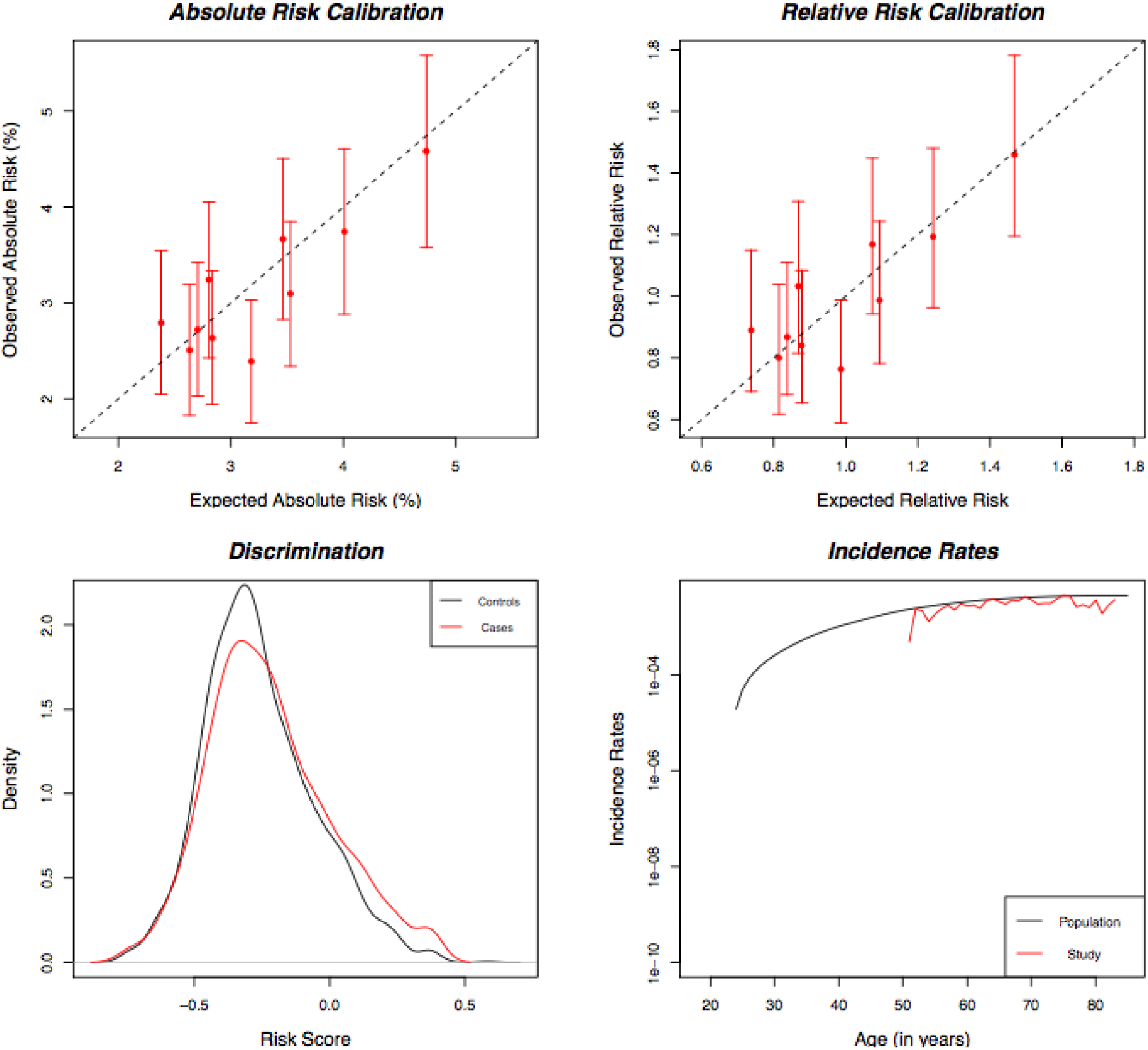
Plots showing model validation results. The top left panel shows calibration results for absolute risk; the top right panel shows calibration for relative risk with respect to the average risk in the validation study; the bottom left panel shows the distribution of risk score in cases (red) and controls (black); the bottom right panel shows the population incidence rates (black) and the incidence rates estimated in validation study (red).

### Additional Options

**iCARE** provides several advanced options as well. For example, `model.ref.dataset.weights` allows the user to optionally specify a vector of weights for each row in the reference dataset. Whenever any averaging is performed over the reference dataset, such as in the case of missing covariates for prediction, a weighted average is applied using the provided sampling weights. Using the `computeAbsoluteRiskSplitInterval` function a user can specify that the absolute risk interval be computed in two parts, using two different sets of parameters. This allows the proportional hazards assumption to be relaxed to some extent, by allowing the relationship between risk factors and the outcome to vary over time. For example, it is well documented that the relationships between certain risk factors, such as body mass index, and breast cancer are different among premenopausal and postmenopausal women. Using `computeAbsoluteRiskSplitInterval`, users can specify a different set of relative risks through the input parameters `model.log.RR` and `model.log.RR2` for use prior to and after a cutpoint of age 50, the median age of menopause. This function is also useful when the distribution of risk factors varies with age.

In addition to returning risk estimates for the specified profiles, the **iCARE** functions can optionally return the absolute risks for the reference dataset as well if `return.refs.risk=TRUE`. The risk scores, or *β*^*T*^ *Z*_*i*_, for the covariate profiles can be obtained by setting the parameter `return.lp=TRUE`. For individuals where there is missing data in covariate profile *Z*, the reported linear predictor is the average of the full linear predictors of all referent subjects in the matching strata according to the approach described in Section 2.5.

## 5. Conclusion

The **iCARE** package is a new tool for building, validating and applying absolute risk models by synthesizing data sources on key model parameters. The tool standardizes methodology and gives researchers the ability to easily update and share absolute risk models, and to evaluate the public health implications of etiologic findings by translating relative risks onto the absolute risk scale. The package incorporates calibration to population-based age-specific disease incidence rates and handling of missing data by leveraging a reference dataset of risk factors for the population of interest. Through its handling of missing data and the ability to incorporate SNP information based on published estimates, the tool gives researchers the ability to easily handle data analytic issues that are likely to arise in practice when building absolute risk models for public health. The package also incorporates standardized methods and tools for validating an absolute risk prediction model in independent studies: a step that is critical for clinical application of risk models (e.g., risk based screening). In this article, we have described the methodology underlying this new tool and illustrated its use with examples by building and validating absolute risk models for breast cancer.

## 6. Appendix

### 6.1 Variance of category specific absolute risk

Here we derive the influence function based variance formula for 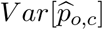 in a nested casecontrol study setting, where 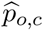 is the observed proportion of cases in the cateogry *c*. We denote by *p*_*c*_, the probability limit of 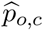. This estimator can be obtained by solving the estimating equation based on the total number of subjects (*N*) in the cohort:

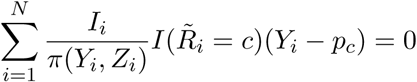

The influence function representation of this estimator is given by:

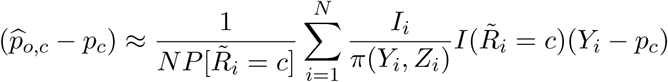

Based on this influence function representation, the asymptotic variance formula is given by:

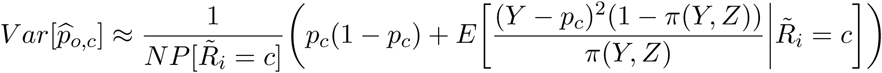

The variance formula includes a binomial variance term and a correction factor that comes in due to the adjustment for non-random sampling of subjects in the nested case-control study. For the full cohort setting, the second term will be zero and only the first term contributes to the 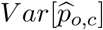

### 6.2 Variance of category specific relative risk

The category specific relative risk is defined as the ratio of the category specific absolute risk and the average absolute risk in the validation study. The categories include approximately equal number of subjects. Hence, the average absolute risk can be approximated by the average of the absolute risk over the categories. We assume that the vector of the centered and scaled category specific absolute risks follows a multivariate normal distribution with mean zero and a diagonal variance-covariance matrix (Σ_*AR*_) with the diagonal entries given by the category specific variance of absolute risk. The vector of category specific relative risks is a smooth function of the vector of category specific absolute risks. Hence we implement a delta method to compute the variance-covariance matrix (Σ_*RR*_) of the vector of category specific relative risks.

We denote the category specific observed relative risk to be 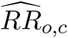 and the category specific predicted relative risk to be 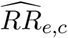. We further denote, 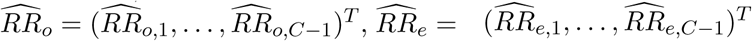 and 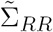 denote the corresponding (*C* − 1) by (*C* − 1) submatrix of Σ_*RR*_. We implement a test for relative risk calibration using the test statistic 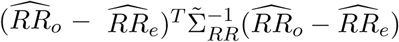 that can be approximated by a chisquare distribution with (*C -* 1) degrees of freedom.

### 6.3 Variance of Area Under the Curve (AUC)

We denote by *I*_1_(*I*_0_), the indicator of inclusion of a case(control) in the cohort study to the nested case-control sample. Let *n*_1_(*n*_0_) be the number of cases (controls) in the cohort. The inverse probability weig ted estimator 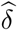 is obtained by solving the estimating equation:

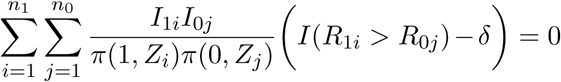. The influence function representation of this estimator is given by:

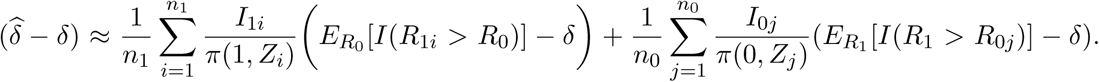

Based on this representation, the asymptotic variance formula is given by:

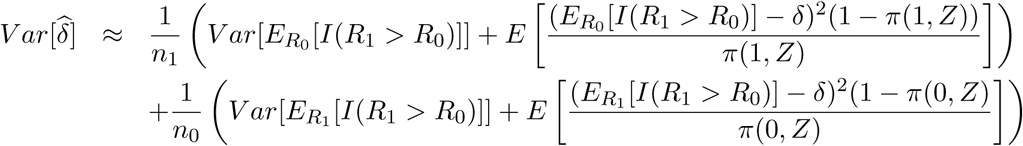

### 6.4 Simulation Study Design for Illustration of Model Validation

We simulate data for a cohort study with 50,000 subjects with information on age of study entry, age of study exit, time to disease onset, disease status, epidemiologic risk factors (i.e., family history and parity) and the 15 breast cancer associated SNPs in `bc_15_snps`. We generate family history from a Bernoulli distribution with success probability 0.14 and parity as a categorical variable taking values 0, 1, 2, 3 with probabilities 0.08, 0.07, 0.25, 0.6, respectively. We generate the 15 breast cancer associated SNPs using the allele frequencies available in `bc_15_snps` and appropriately accounting for correlation with family history by the methods described in Section 2.6. We use the log relative risks for family history and parity given in `bc_model_log_or`. For the SNPs, we use the log odds ratios given in `bc_15_snps`. We generate an age of disease onset from a Cox model that assumes multiplicative association of the risk factors on disease risk. The baseline hazard function of the Cox model is generated from a Weibull distribution with parameters chosen such that the probability of developing the disease by age 50 and 70 years are approximately 5% and 12% respectively. We generate age of study entry from a discrete uniform distribution in the range 50 years to 72 years. We also generate a length of observed follow-up from a discrete uniform distribution in the range 5 years to 13 years.

After we generate the full cohort data, we select a case-control sample nested within the cohort and assume that data on the 15 SNPs are available only on this sub-sample. We assume a logistic regression model of selection depending on case/control status, age of study entry and observed follow-up and compute the selection probabilities using the following code:

~~~
selection.model.params = c(−7.52,9.47,0.05,0.11,−0.09,0.04)
selection.model.matrix = model.matrix(˜observed.outcome * (study.entry.age + observed.followup), data = validation.cohort.data)
selection.exp.lin.pred = exp(selection.model.matrix %*%selection.model.params)
selection.prob = (selection.exp.lin.pred/(1 + selection.exp.lin.pred))
~~~

In the above code, the `selection.model.params` object gives the log-odds ratios associated with the covariates in the selection model. The selection probabilities are used to generate a binary inclusion (yes = 1, no = 0) variable from a Bernoulli distribution. All the subjects with inclusion taking value 1 are included in the nested case-control sample. Both the full cohort and nested case-control datasets have been made available with **iCARE** under the names `validation.cohort.data` and `validation.nested.case.control.data`, respectively.

## Acknowledgements

The works of Parichoy Pal Choudhury and Nilanjan Chatterjee were supported by the PatientCentered Outcomes Research Institute (PCORI) Award (ME-1602-34530). The works of Paige Maas, Amber Wilcox, David Check and Montse Garcia-Closas were supported by the Intramural Research Program, Division of Cancer Epidemiology and Genetics, National Cancer Institute, National Institutes of Health, Department of Health and Human Services. The statements and opinions in this article are solely the responsibility of the authors and do not necessarily represent the views of the Patient-Centered Outcomes Research Institute (PCORI), its Board of Governors, or Methodology Committee or the National Cancer Institute, National Institutes of Health, Department of Health and Human Services. Conflict of Interest: None declared.

*Journal of Statistical Software* http://www.jstatsoft.org/

published by the American Statistical Association http://www.amstat.org/

Volume VV, Issue II *Submitted:* yyyy-mm-dd

MMMMMM YYYY *Accepted:* yyyy-mm-dd

